# Genetic composition and evolution of the prevalent *Mycobacterium* tuberculosis lineages 2 and 4 in the Chinese and Zhejiang Province populations

**DOI:** 10.1101/2020.11.07.372573

**Authors:** Beibei Wu, Wenlong Zhu, Yue Wang, Qi Wang, Lin Zhou, Zhengwei Liu, Lijun Bi, Mathema Barun, Barry N. Kreiswirth, Liang Chen, Songhua Chen, Xiaomeng Wang, Weibing Wang

## Abstract

The causative agent of tuberculosis (TB) comprises seven human-adapted lineages. Human movements and host genetics are crucial to TB dissemination. We analyzed whole-genome sequencing data for a countrywide collection of 1154 isolates and a provincial collection of 1296 isolates, constructed the best-scoring maximum likelihood phylogenetic tree, conducted Bayesian evolutionary analysis to compute the most recent common ancestors of lineages 2 and 4, and assessed the antigenic diversity in human T cell epitopes by calculating pairwise dN/dS ratios. Of the 1296 Zhejiang isolates, 964 (74.38%) belonged to lineage 2 and 332 (25.62%) belonged to lineage 4. L2.2 is the most ancient sub-lineage in Zhejiang, first appearing approximately 6897 years ago (95% HDI: 6513-7298). L4.4 is the most modern sub-lineage, first appearing approximately 2217 years ago (95% HDI: 1864-2581). The dN/dS ratios revealed that the epitope and non-epitope regions of lineage 2 strains were significantly (*P*<0.001) more conserved than those of lineage 4. An increase in the frequency of lineage 4 may reflect its successful transmission over the last 20 years. The recent common ancestors and transmission routes of the sub-lineages are related to the entry of humans into China and Zhejiang Province.

## Introduction

The causative agent of tuberculosis (TB), *Mycobacterium* tuberculosis (Mtb), is an obligate pathogen that comprises seven human-adapted lineages (Coscolla and Gagneux 2014). Mtb is one of the most successful human pathogens, having killed an estimated 1 billion people over the last 200 years (Gagneux 2012). In 2017, TB caused an estimated 1.6 million deaths, including 300,000 deaths in the HIV-positive population. Sustained reductions in disease incidence of up to 20% per year are required to meet the targets set out in the “WHO END TB” Strategy (Glaziou et al. 2013; Leung et al. 2018). However, current estimates suggest that the incidence is decreasing at a rate of only 1.5% per annum (WHO 2018).

It is well known that the social characteristics of human populations (Lonnroth et al. 2009), host genetics (Gagneux 2012) and human interventions (e.g., the implementation of disease control programs) are crucial determinants of TB. Accumulating evidence indicates that human migrations and activities influence the population structure of Mtb (Nathanson et al. 2010). As such, human-adapted Mtb lineages have shown a strong phylogeographic population structure in which different lineages are associated with distinct geographic regions (Filliol et al. 2006; Hershberg et al. 2008; Reed et al. 2009). A number of studies have found differences in virulence and immunogenicity among the seven lineages (Coscolla and Gagneux 2010; Parwati et al. 2010). Interestingly, the extent of their geographic distribution differs markedly, with some exhibiting a global distribution while others showing a strong geographic restriction. Widely distributed Mtb is more likely to spread. Therefore, identifying the predominant lineages in various regions can provide critical insight into the successful transmission and development of TB.

The human-adapted members of *Mycobacterium tuberculosis complex* (MTBC) can be classified into seven independent lineages (Coscolla and Gagneux 2014), all of which have humans as their only known host. Lineages 2 and 4 appear to be more virulent and transmissible on average than the other Mtb lineages (Coscolla and Gagneux 2014; Liu et al. 2018). However, this is not always true, and there is a great deal of variation among the lineage 4 strains. Lineage 2, which is also known as the East-Asian lineage due to its predominance in East Asia, includes the Beijing family of strains that have received particular attention because they are associated with drug resistance and virulence and are considered to be a ‘successful’ lineage (Nathanson et al. 2010). Molecular epidemiological studies have reported considerable variation in the transmission success of lineage 2 strains. For example, several studies using whole-genome sequencing (WGS) have demonstrated that lineage 4 can be further subdivided into several sub-lineages (Coll et al. 2014; Stucki et al. 2016). These sub-lineages partially reflected strain families that had been previously defined based on various genotyping techniques. The increase in human population density during the agricultural and industrial revolutions would then have selected for increased virulence in some Mtb lineages.

To understand the phenotypic consequence of between and within lineage diversity, one can look at its evolutionary conservation of protein residue (Shih et al. 2012), as between-lineage differences in the sharing of mutations may impact their phenotypes. Between-strain comparison of genomic regions encoding proteins that are recognized by human T cells has revealed that T cell epitopes are among the most conserved regions in the Mtb genomes; they exhibit lower frequencies of amino acid changes compared to essential genes and non-epitope antigen regions (Coscolla et al. 2015; Yruela et al. 2016).

It remains unclear when epidemic forms of TB first arose in China, how the strains transmitted successfully within China, and what course these epidemics may have followed throughout Chinese history. In the present study, we reconstruct the phylogenomic history of epidemic TB in eastern China and use it to examine how the intersection of Mtb phylogeny, geography and demography has contribute to the widespread dispersal of TB in this country. We examine the SNPs (single nucleotide polymorphisms) shared by the predominant lineages in China as a means to explore the common genetic characteristics that have contributed to its wide transmission. Our analyses provide insights into the genomic polymorphism of the predominant TB lineages and the genetic basis for the widespread dissemination capacity and virulence of this important human disease.

## Results

### Collection and genomic sequencing of1296 Mtb isolates from Zhejiang Province

From 1998 to 2013, a total of 1434 clinical isolates were collected; of them, 1372 (95.67%) were culture-positive and 1329 (96.87%) met our predefined criteria for the sequencing purity and concentration. Thirty-three isolates that were cross-contaminated or did not represent Mtb were excluded. In total, 1296 isolates were included for our analysis (Figure 1).

**Figure 1.**
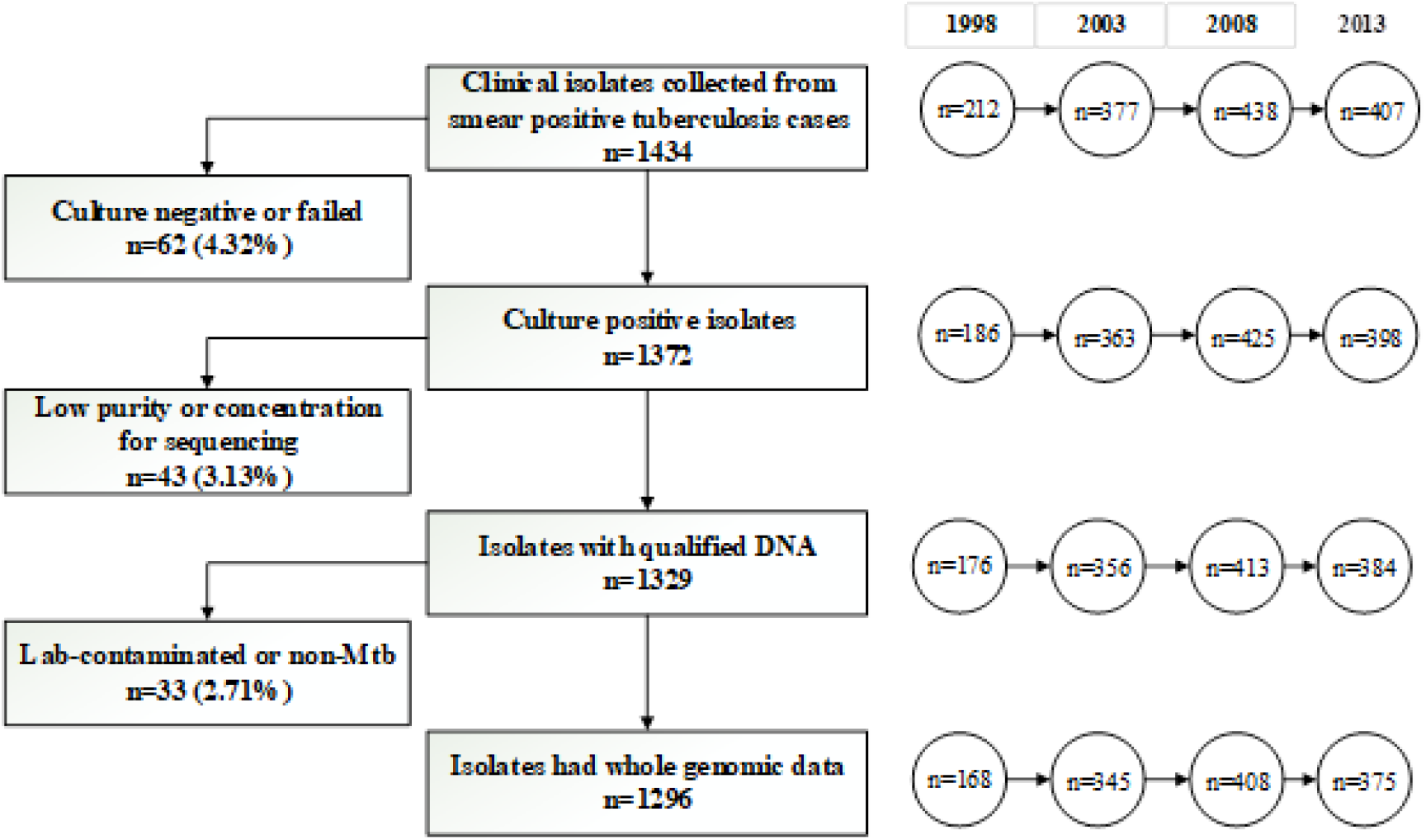
Clinical isolates collected in Zhejiang, 1998-2013.

### Phylogenetic characteristics of the lineage 2 and lineage 4 strains

WGS data were obtained from the 1296 Mtb isolates from Zhejiang Province and downloaded for 1154 previously studied isolates that were obtained from around the world and represented the six main previously-defined phylogeographic lineages of Mtb. These data were used to construct phylogenetic trees (Figure 2). Of the 1296 Zhejiang isolates, 964 (74.38%) belonged to lineage 2 and 332 (25.62%) belonged to lineage 4. We next selected a subset of lineage 4 clinical isolates (n=771) from 17 countries and a subset of lineage 2 clinical isolates (n=383) from 12 countries. To determine the placement of the Zhejiang strains along the evolutionary path of these lineages, we reconstructed maximum-likelihood phylogenies for lineages 2 and 4. The phylogenetic trees showed that lineage 2 comprises three sub-lineages, L2.1 (10.17%), L2.2 (32.57%) and L2.3 (57.26%); among them, L2.3 (552 strains) was the predominant sub-lineage in Zhejiang Province, accounting for 42.59% of the total strains. Lineage 4 was found to comprise three sub-lineages, L4.2 (18.07%), L4.4 (38.56%) and L4.5 (43.37%).

**Figure 2.**
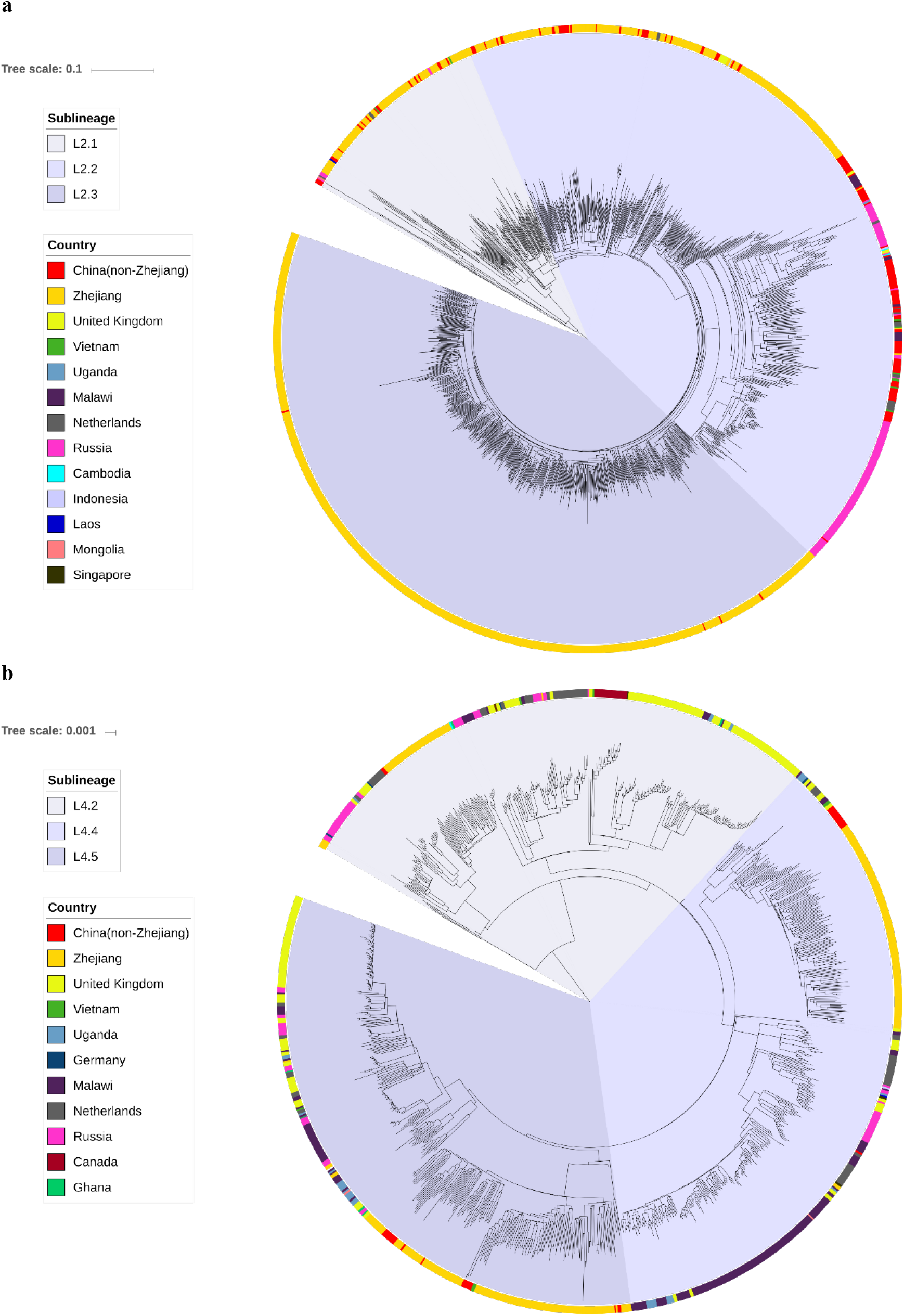
Bayesian phylogeny of the Zhejiang *M. tuberculosis* isolates and 1154 globally distributed publically available genomes for (a) lineage 2 and (b) lineage 4. Scale bar indicates the regions of origin. The *M. tuberculosis* sub-lineages, L2.1, L2.2, L2.3, L4.2, L4.4 and L4.5, are indicated respectively.

The distributions of sub-lineages varied between the administrative/geographic regions of Zhejiang Province (East, North, West, South and Middle). The lineage 4 types accounted for the largest proportion in Southern Zhejiang (40.10%), while Western Zhejiang had the lowest proportion (19.57%) of these lineages. Analysis of spatial-temporal trends in the distributions of lineage 2 and 4 isolates among the five districts indicated that the proportion of lineage 4 isolates decreased in Northern and Southern Zhejiang over the 16-year study period, whereas it increased in Western Zhejiang (Supplementary_Fig_S1.).

### Phylogeographic evolution of the major sub-lineages

Published phylogeographic studies have indicated an African origin for Mtb, suggesting that it was introduced to other continents via human migration (Hershberg et al. 2008; Comas et al. 2013). To further explore the evolutionary relationship of these strains and their geographical distribution, we used Bayesian evolutionary analysis (Table 1, Figure 3) to predict the divergence time of the most recent common ancestors of four sub-lineages (Supplementary_Fig_S2.).

**Table 1.**
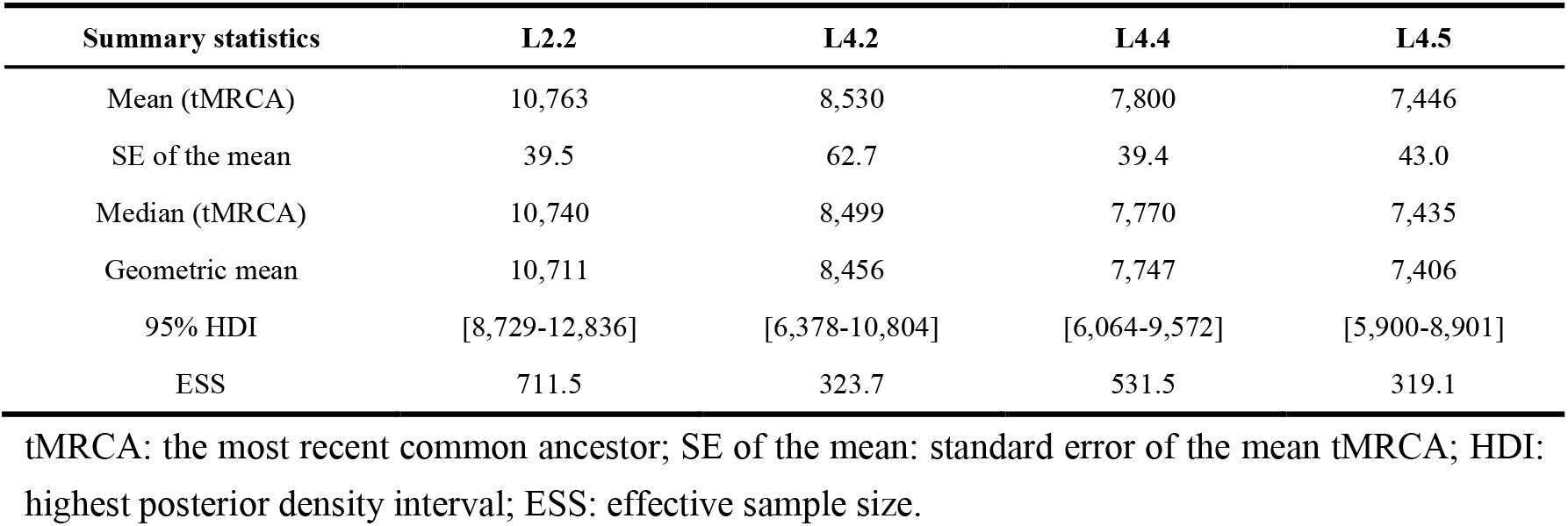
The most recent common ancestors of L2 and L4 sub-lineages in China

| Summary statistics | L2.2 | L4.2 | L4.4 | L4.5 |
| --- | --- | --- | --- | --- |
| Mean (tMRCA) | 10,763 | 8,530 | 7,800 | 7,446 |
| SE of the mean | 39.5 | 62.7 | 39.4 | 43.0 |
| Median (tMRCA) | 10,740 | 8,499 | 7,770 | 7,435 |
| Geometric mean | 10,711 | 8,456 | 7,747 | 7,406 |
| 95% HDI | [8,729-12,836] | [6,378-10,804] | [6,064-9,572] | [5,900-8,901] |
| ESS | 711.5 | 323.7 | 531.5 | 319.1 |

tMRCA: the most recent common ancestor; SE of the mean: standard error of the mean tMRCA; HDI: highest posterior density interval; ESS: effective sample size.

**Figure 3.**
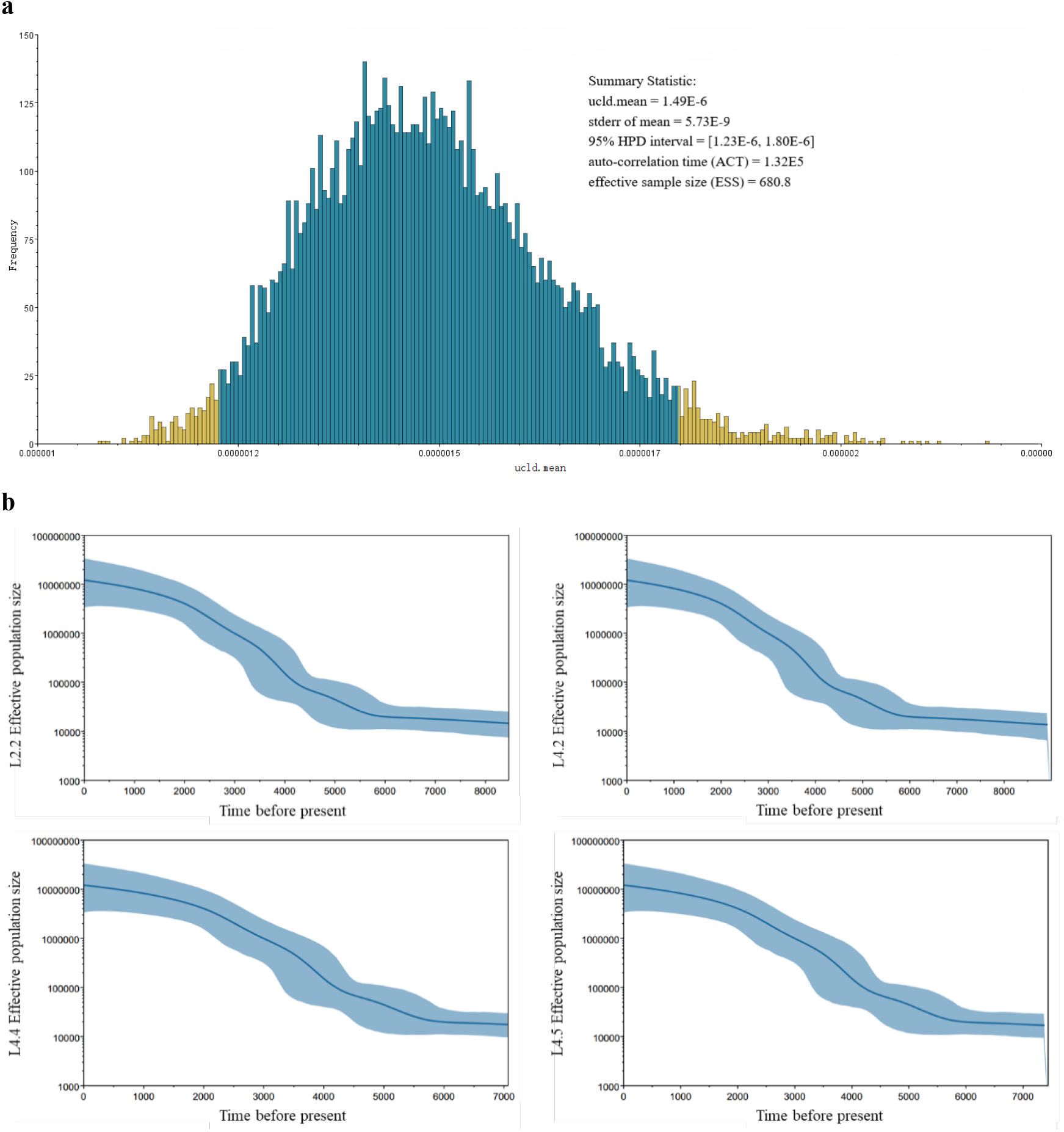
Mutation rates and changes in sub-lineage diversity over time. (a) The mutation rate was estimated using Beast. (b) Bayesian skyline plots indicating changes in the diversity of four sub-lineages over time. Shadowed areas show the 95% HPD (high posterior density) intervals for the population-size estimations.

Our results revealed that L2.2 is the most ancient of the studied sub-lineages in China, with its tMRCA appearing around 10,763 years ago (95% HDI: 8729-12,836 years ago), whereas L4.5 is the most modern of the studied sub-lineages in China, with its tMRCA appearing around 7446 years ago (95% HDI: 5900-8901). As shown in Figure 3a, the substitution rate of Mtb was found to be a mean of 4.35×10^−9^ substitutions per genome per site per year [95% HPD interval: 3.58×10^−9^-5.26×10^−9^; converted by the calculated annual mutation rate of each polymorphic locus (24,633 loci): *ucld.mean*=1.49×10^−6^].

Given the times of origin for the four sub-lineages in China, the characteristics of the MCC tree (Supplementary_Fig_S2.), and historical information on the arrival and spread of modern humans in China (Comas et al. 2013), we propose two possible routes of propagation across China for each of the studied sub-lineages (Figure 4). For L2.2, one potential route of propagation originates in Xinjiang in Northwest China and spreads to the South and Southeast, while the other originates in Fujian and spreads to the north. For L4.2, one potential route of propagation originates in Qinghai Province in Western China and spreads to the East and Southeast, while the other originates in Heilongjiang Province in Northeast China and spreads to the South. For L4.4, one possible route of propagation originates in Guangdong and Hunan Provinces of Southern China and spreads to the North, while the other originates in Heilongjiang Province and spreads to the South. For L4.5, one possible route of propagation originates in Xinjiang Province and spreads to the East and Southeast, while the other originates in Heilongjiang Province and spreads to the South and Southwest. The origin times of some key propagation points are shown in Figure 4.

**Figure 4.**
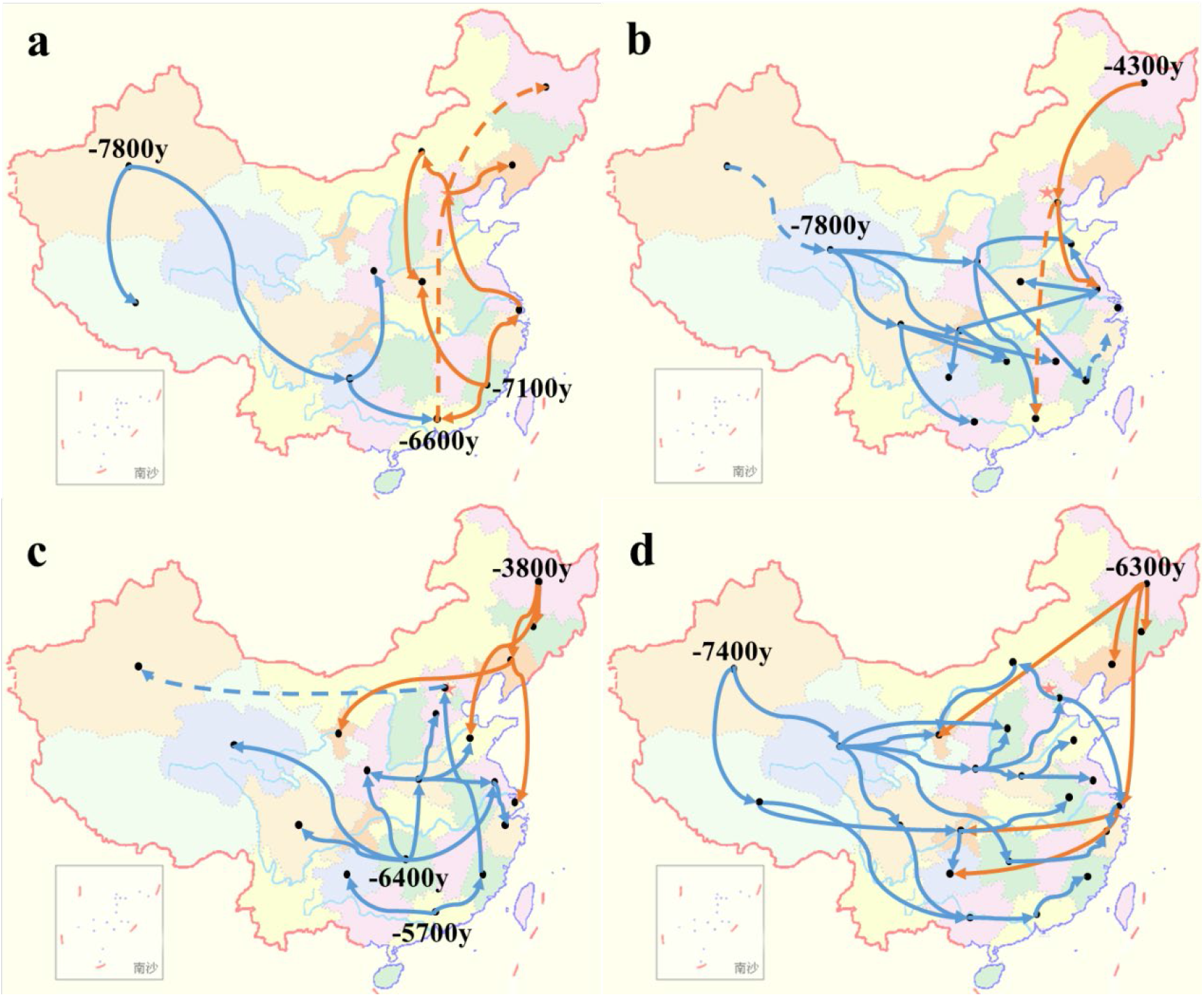
Potential propagation routes of four sub-lineages in China. Shown are routes for L2.2 (a), L4.2 (b), L4.4 (c) and L4.5 (d). The dotted line indicates that the distance is long and the evidence maybe weak (possibly due to a lack of strains). Blue lines indicate older transmission routes, while orange lines indicate more recent transmission routes.

We used a similar method to obtain the divergence times for the MRCAs of the six sub-lineages found in Zhejiang Province. As shown in Table 2, we found that L2.2 is the most ancient of the studied sub-lineages in Zhejiang, with its MRCA appearing around 6 897 years ago (95% HDI: 6513-7298 years), while L4.4 is the most modern of the studied sub-lineages in Zhejiang, with its MRCA appearing around 2217 years ago (95% HDI: 1864-2581 years).

**Table 2.**
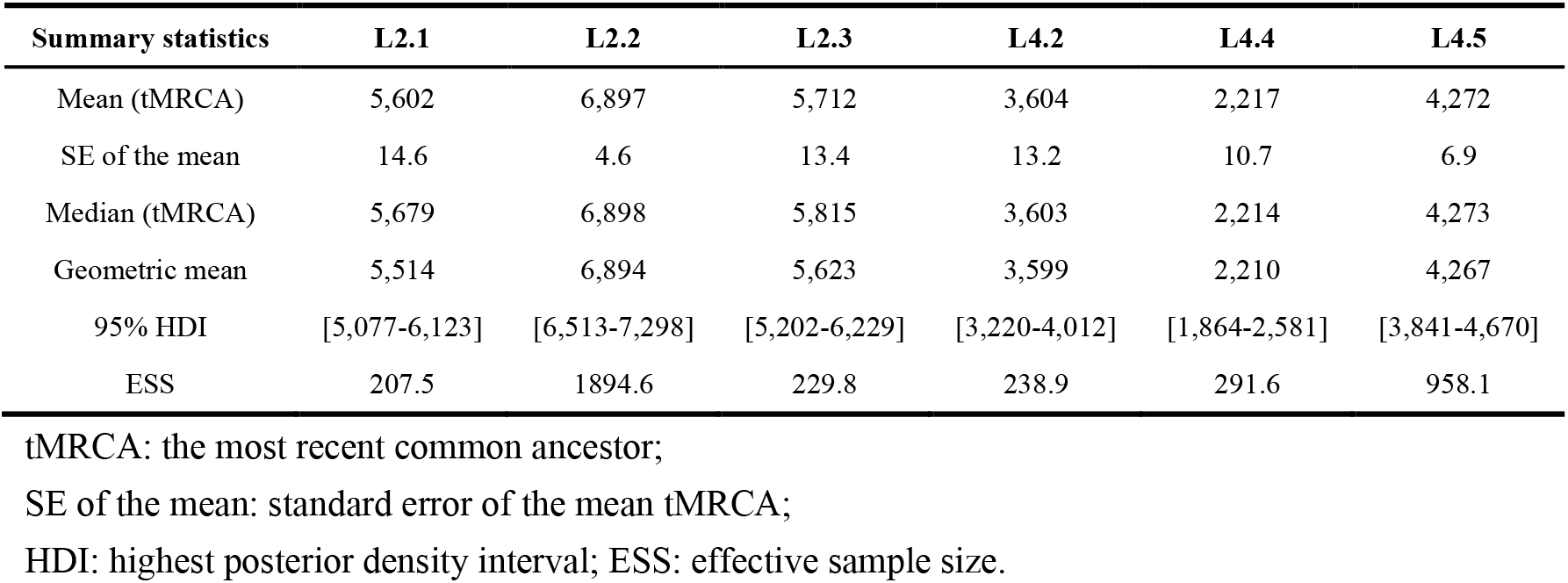
The most recent common ancestor of L2 and L4 in Zhejiangx

| Summary statistics | L2.1 | L2.2 | L2.3 | L4.2 | L4.4 | L4.5 |
| --- | --- | --- | --- | --- | --- | --- |
| Mean (tMRCA) | 5,602 | 6,897 | 5,712 | 3,604 | 2,217 | 4,272 |
| SE of the mean | 14.6 | 4.6 | 13.4 | 13.2 | 10.7 | 6.9 |
| Median (tMRCA) | 5,679 | 6,898 | 5,815 | 3,603 | 2,214 | 4,273 |
| Geometric mean | 5,514 | 6,894 | 5,623 | 3,599 | 2,210 | 4,267 |
| 95% HDI | [5,077-6,123] | [6,513-7,298] | [5,202-6,229] | [3,220-4,012] | [1,864-2,581] | [3,841-4,670] |
| ESS | 207.5 | 1894.6 | 229.8 | 238.9 | 291.6 | 958.1 |

tMRCA: the most recent common ancestor; SE of the mean: standard error of the mean tMRCA; HDI: highest posterior density interval; ESS: effective sample size.

Given the origin times of the six sub-lineages in Zhejiang, the characteristics of the MCC tree (Supplementary_Fig_S3.) and the above-described possible transmission routes of the four sub-lineages in China, we inferred the potential propagation routes for the six sub-lineages in Zhejiang, as shown in Figure 5. The directions and estimated years at which the strains entered Zhejiang from other regions are basically consistent with the transmission routes of the four sub-lineages (L2.2, L4.2, L4.4 and L4.5) in China. More specifically, L2.1 and L2.3, which derived from 5,700 years ago, might be related to the origin and migration of Liangzhu Culture (about 5,500 years ago), sharing similar original time and geographical distribution (Yi 2019). L4.5, deriving from 3,600 years ago, might be related to the Battle of Mingtiao, which was the final battle of the Xia Dynasty (circa 1,600 BC). Shang Tang won the battle and Xia Jie retreated to Nanchao, adjacent to Zhejiang Province (Fan 2017). L4.4, deriving from 2,200 years ago, might be related to the war of Qin State destroying Chu State (circa 200 BC). At that time, the territory of Chu included western and southeastern Henan, southern Shandong, Hubei, Hunan, Jiangxi, Anhui, Jiangsu, and Zhejiang. The marching route of Qin destroying Chu was consistent with the transmission route of L4.4 (Li 1981). Moreover, the spread of L2.2 might be related to the origin of Zhejiang’s agricultural civilization and the transmission route of L4.5 began from sea, which may be related to the origin of the Maritime Silk Road (CCTV 2007).

**Figure 5.**
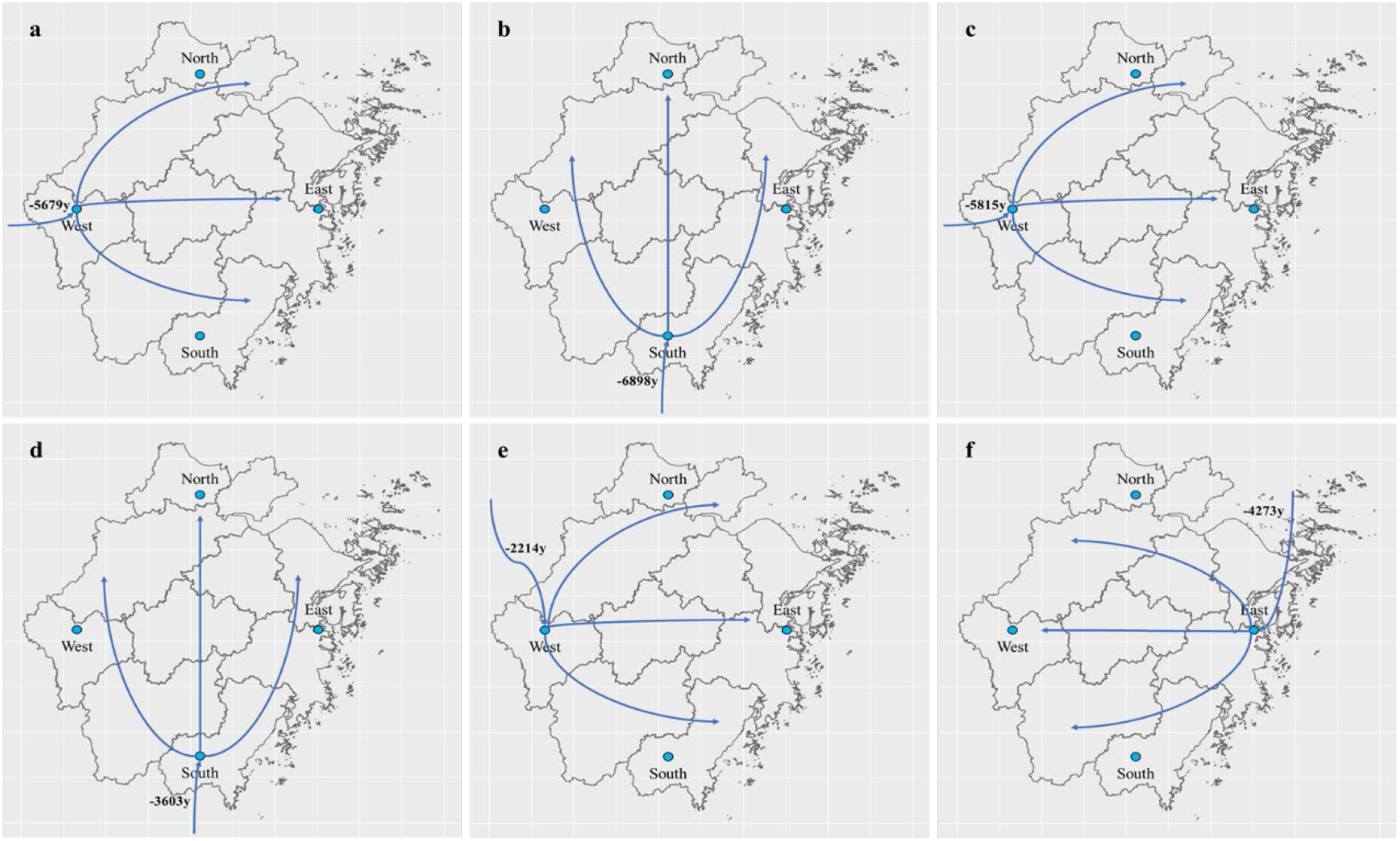
Potential propagation routes of six sub-lineages in Zhejiang Province. Shown are L2.1 (a), L2.2 (b), L2.3 (c), L4.2 (d), L4.4 (e) and L4.5 (f). The curves without starting points indicate the directions and years of the strains entering Zhejiang Province from other regions.

### Genomic features of lineages 2 and 4

We compared the genetic diversity of the lineage 2 and 4 strains in Zhejiang Province to that of the global strains. As seen in the global strains, there was greater genetic diversity among the lineage 4 strains from Zhejiang Province than among the lineage 2 strains (Figure 6). Zhejiang lineage 4 strains harbored a mean diversity of 565 SNPs between any two strains, compared to 291 SNPs in lineage 2.

**Figure 6.**
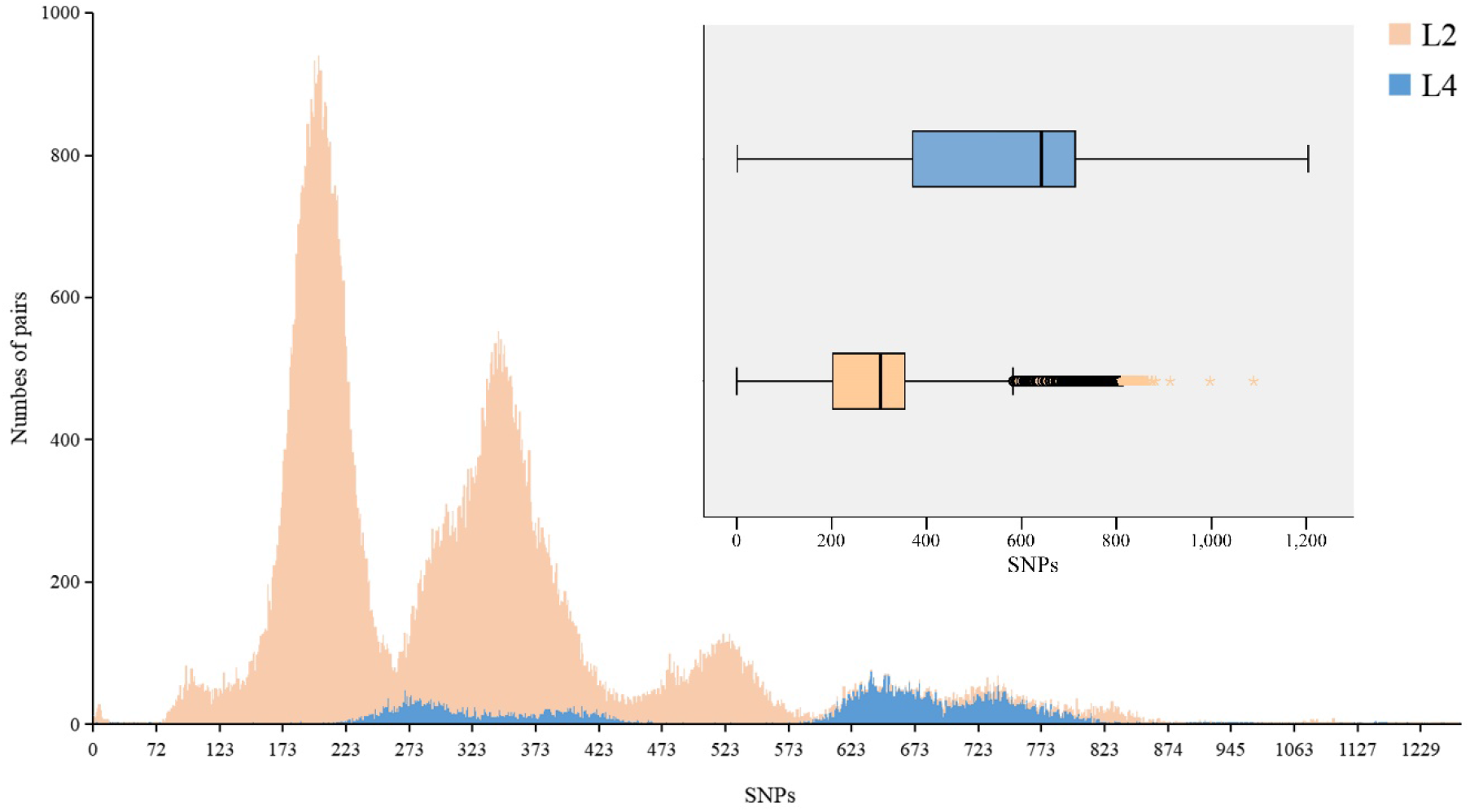
Number of pairwise differences between Mtb strains for lineage 2 and lineage 4. The alignment of 217 human-adapted Mtb clinical strains published previously (Comas et al., 2013) was used to calculate pairwise differences of global strains.

Our estimation of the genetic diversity among the sub-lineages of lineages 2 and 4 based on the SNP pairwise distances showed that L2.3, the predominant sub-lineage in lineage 2, was significantly more conserved than L2.1 (mean of 202 and 337 SNPs, respectively, shared between isolate pairs; Wilcoxon rank-sum test, P < 0.0001). In lineage 4, we observed the opposite trend, as the predominant sub-lineage, L4.5, was more diverse than L4.2 (mean of 385 and 253 SNPs, respectively; Wilcoxon rank-sum test, P < 0.0001) (Figure 7).

**Figure 7.**
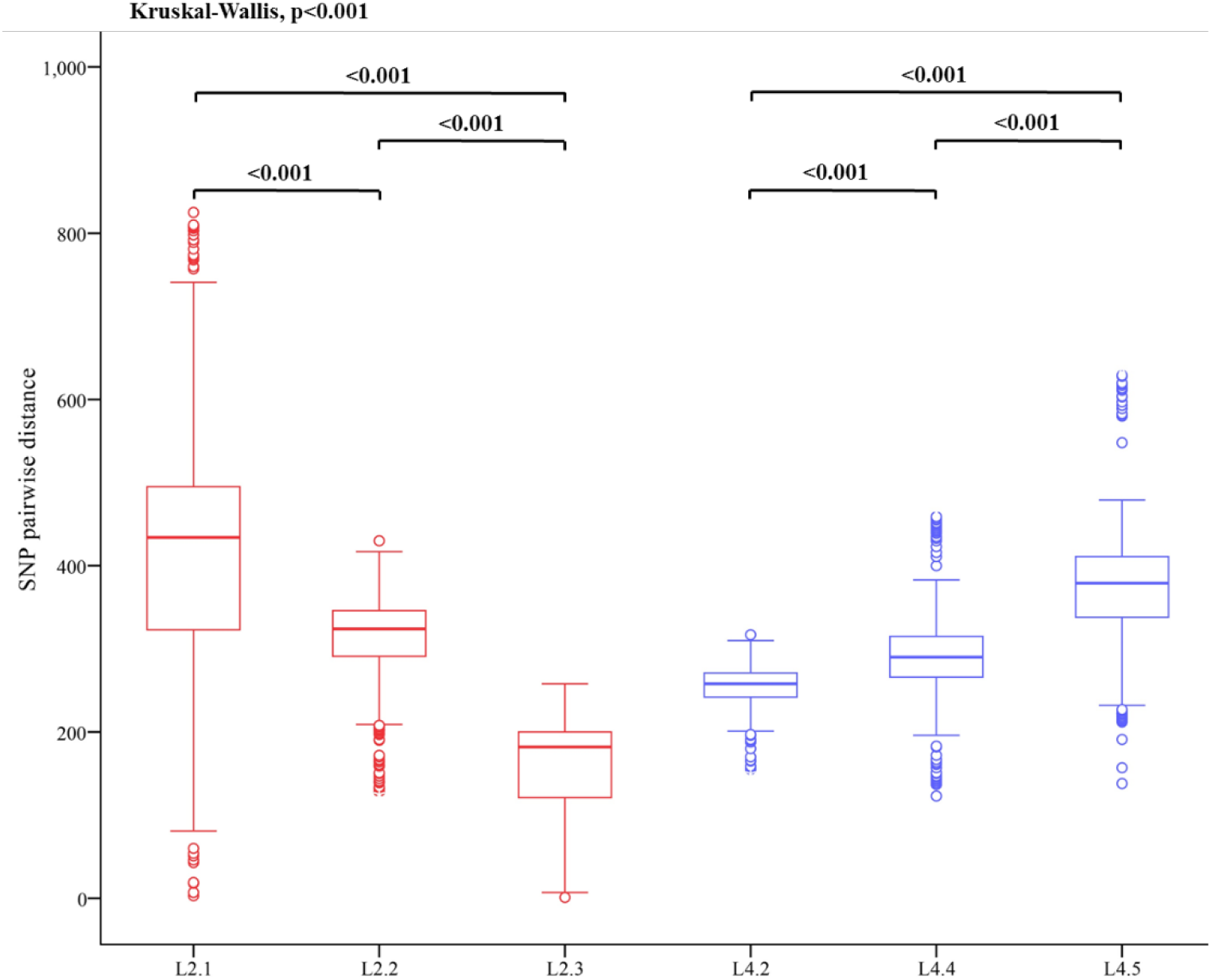
Box plots of pairwise genetic distances (number of polymorphisms) for each sub-lineage.

To assess the genetic diversity of antigens in the lineage 2 and 4 strains, we calculated the non-synonymous to synonymous substitution (dN/dS) ratios for the epitope and non-epitope regions, along with the distribution of amino acid replacements in individual epitopes. We found that the dN/dS ratio of epitope and non-epitope regions exhibited significantly more conservation in lineage 2 strains than in lineage 4 strains. In lineage 2 strains, however, the T cell epitope regions showed significantly higher dN/dS ratios than the non-epitope regions (Figure 8). When we assessed the evolutionary conservation of human T cell epitopes in the sub-lineages of lineage 2 and lineage 4 (Supplementary_Fig_S4.), we found that the dN/dS ratios for the sub-lineages of lineage 4 were similar to those of the overall lineage. For the sub-lineages of lineage 2, meanwhile, the dN/dS ratio of the lowest-prevalence sub-lineage, L2.1, differed from that of the overall lineage, whereas the ratios of the other sub-lineages were consistent with those of the overall lineage.

**Figure 8.**
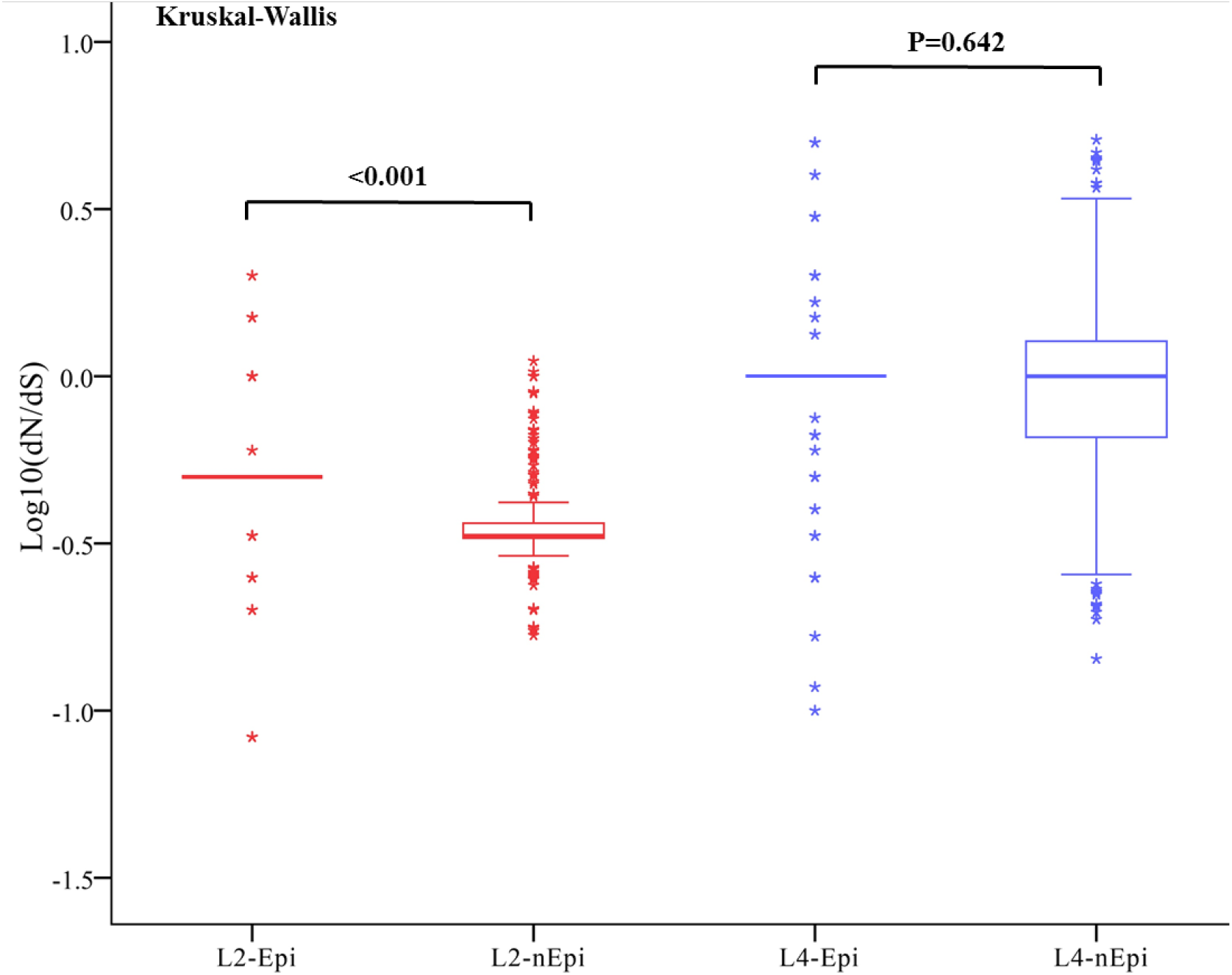
Pairwise ratios for the rates of nonsynonymous to synonymous substitutions (dN/dS) in lineage 2 and 4 isolates, assessing epitope and non-epitope regions of T cell antigens.

When we analyzed the distribution of amino acid replacements in individual epitopes, we found that a large majority (95%) of the 491 T cell epitopes showed no amino acid change (Supplementary_Fig_S5.). However, lineage 2 had more epitopes that harbored at least one amino acid change, compared to lineage 4. In lineage 2, four epitopes (*esxL, lpqH, fbpB* and *lppX*) harbored more than two variable positions.

## Discussion

Whole-genome sequencing of 1296 Zhejiang Province strains and comparison with 1154 publically-available global MTBC genomes was used to elucidate the distribution of MTBC sub-lineages in the Chinese population. Genetic diversity and T cell epitopes were significantly different between sub-lineages.

We observed differences in the spatiotemporal characteristics of the lineage 2 and lineage 4 strains. While the proportion of lineage 4 strains in Western Zhejiang was generally low, the proportion of cases arising from lineage 4 strains increased over time across the four survey periods. This increase may reflect the successful transmission of these strains over time. Other studies in various settings have reported that the higher fitness of lineage 2/ Beijing strains is reflected by increases in their frequency over time (Tuite et al. 2013). In contrast, the frequency of lineage 4 strains in Southern Zhejiang showed a downward trend, which is incompatible with the above hypothesis.

A previous study showed that migrants had an impact on the spread of Mtb in Russia (Mokrousov 2013). Lineage 4 was found at a high proportion in the Southern Zhejiang, which is typically the destination choice of migrant population from other provinces. Relatively low migration has been seen in the Western region of Zhejiang Province; however, due to developments in the economy and convenience of transportation, migration into this region increased significantly between 2000 and 2010. The similarity between the characteristics of migration and the trends in the proportion of lineage 4 suggest that there is likely to be a relationship between lineage 4 and migration. Future studies will be needed to assess whether migrants increase the risk of lineage 4 transmission in Zhejiang. Our Bayesian evolutionary analyses suggest that the identified sub-populations of Mtb emerged in China around 1000 years ago, expanded in parallel from the 12^th^ century onwards, and peaked (at a whole-population level) in the late 18^th^ century. More recently, sub-lineage L2.3, which is indigenous to China and exhibits relatively high transmissibility and extensive global dissemination, came to dominate the population dynamics of Mtb in China.(Liu et al. 2018)

The tMRCAs that our Bayesian evolution model calculated for the four sub-lineages are related to the entry of modern humans into China, their migration routes, and the expansion of the population in the Neolithic Age (about 10,000 years ago). We found that the population sizes all four sub-lineages increased significantly around 5000 years ago, which coincides with the origin of the Chinese civilization according to the historical record (Comas et al. 2013). During that period, the population grew on a large scale and engaged in frequent social activities, presumably accelerating the evolution and spread of Mtb.

The Mtb strains differ genetically in their content of SNPs, and the more recently transmitted strains would be expected to have reduced levels of genetic diversity. Our findings show that number of pairwise differences between Mtb strains for lineage 2 in Zhejiang province was lower than that in global strains, whereas the opposite is true for lineage 4. The strains of lineage 2, which represent the predominant clades in Zhejiang, are separated by a smaller genetic distance, indicating more ongoing transmission. In contrast, the lineage 4 strains may be more likely to represent external inputs. The sub-lineages also differ in their genetic diversity, with sub-lineage L2.3 (the predominant within lineage 2) showing lower genetic distances compared to L2.1 and L2.2. Therefore, our results suggest this discrepancy supports the idea that there is an epidemiologic distinction between lineage 2 and lineage 4 in Zhejiang Province.

We detected three main potential routes for the spread of MTBC: the first originates in Xinjiang (about 8000 years ago) and may be traced back to human migration through the Eurasian continent from Europe to Central Asia, and then to East Asia (beginning around 15,000-18,000 years ago) (Zhong et al. 2011); the second is consistent with the initial arrival of modern humans in South and Southeast Asia, followed by their entry into China by sea ~ 8000 years ago (Gray and Jordan 2000; Barton et al. 2009) and their subsequent spread to Southeastern China (Fujian, Guangdong and Hunan) about 6000 years ago; and the third and most modern route originates in Heilongjiang (3000-6000 years ago) and may trace back to Japan and Korea. These results are consistent with those of a previous study (Comas et al. 2013). Our findings also support the idea that MTBC is a very old bacterium whose spread in China was achieved through the entry of modern humans into the country and their subsequent expansion and development of agricultural civilization (8000 years ago) (Wirth et al. 2008).

The substitution rate per site per year obtained in our study was essentially the same as the genomic-level prediction (2.58×10^−9^, 95% HPD interval: 1.66×10^−9^ to 2.89×10^−9^) obtained by Comas et al. (Comas et al. 2013). However, this rate is much lower than recent estimates of short-term substitution rates for experimental models of TB and human outbreaks of the disease (Ford et al. 2011; Walker et al. 2013). Deleterious mutations tend to disappear during long-term evolution due to purifying selection, while the substitution rates tend to increase in experimental strains due to positive selection. This may explain why the substitution rate for long-term evolution is much lower than the short-term substitution rate.

We hypothesized that lineages that are predominant in a specific human population and undergoing ongoing transmission have a higher fitness and virulence (Rodrigo et al. 1997; Ernst 2012). In our study, as expected, essential genes were more conserved than nonessential genes, and a large majority of the currently known T cell antigens were completely conserved, in agreement with previous reports for the Mtb overall (Comas et al. 2010; Coscolla et al. 2015). TB does not use antigenic variation as a main mechanism of immune evasion, and other studies found that reduced and/or delayed inflammatory responses were associated with increased Mtb virulence (Tsenova et al. 2005; Subbian et al. 2013). However, for both predominant lineage 2 and predominant sub-lineage L2.3, we obtained significantly higher dN/dS ratios for the T cell epitopes compared to the non-epitope regions. Other studies had found that although the majority of human T cell epitopes in Mtb were conserved (Comas et al. 2010) and relatively few of its antigens and epitopes exhibit evidence of diversifying selection and antigenic variation, the diverse regions exhibit nucleotide diversities and dN/dS ratios higher than the genome-wide average (Oleksyk et al. 2010). We identified four antigens that exhibited more than two nonsynonymous variations in the epitope regions of both lineages: *esxL, lpqH, fbpB* and *lppX*. Notably, these sites also exhibited diversity across the different successful sub-lineages. This natural sequence diversity suggests that variation in these particular antigens might benefit the pathogen, such as by allowing it to escape from human T cell recognition. Future studies will be needed to assess how the limited diversity in Mtb T cell epitopes can impact immune escape. It is the conversation of most T cell epitopes in lineage 2 stains making them the delayed inflammatory immune response and increased virulence at a later stage, meanwhile, the diversity of some epitopes made them affect a wider population.

In conclusion, our study indicates that the spatiotemporal distribution characteristics of lineage 2 and 4 strains in Zhejiang Province are changing and the increase in the frequency of lineage 4 may reflect its successful transmission over the last 20 years. We reconstruct the phylogenomic history of TB transmission and analyze genomic features of lineages 2 and 4 in order to understand the intersection of phylogeny, geography, and demography to gain some insights about TB epidemics.

## Materials and Methods

### Study population and samples

The study population included patients with pulmonary disease and culture-positive TB sampled from 12 locations in Zhejiang Province of Eastern China during drug-resistance surveillances performed in 1998, 2003, 2008 and 2013. The same protocol was applied in all four surveillance periods. For each of the 12 locations, we randomly enrolled 30 new smear-positive patients and all previously treated smear-positive patients.

New cases were defined as those who had never received TB drugs or who had received treatment for less than 1 month. Previously treated cases were defined as those who had received previous TB treatment for 1 month or longer. All patients were active TB cases with bacteriological confirmation by sputum culture. Newly diagnosed patients provided three sputum specimens (spot, morning, and night) and previously treated patients provided two sputum specimens (spot and morning or night). Epidemiological data were collected by trained doctors at TB-designated hospitals, and patients were surveyed on site using a standard questionnaire. Demographic data for the study population are provided in Supplementary_Table_S1.

Samples were tested for Mtb by microscopy and culture in a manner consistent with national guidelines (China 2017). Isolates were cultured on Middlebrook medium for 4-6 weeks at 37°C and DNA was extracted using magnetic beads (Tiangen Biotech Co., Ltd.). Rifampicin and isoniazid drug-susceptibility testing was performed using the proportion method in Löwenstein-Jensen medium (Aziz et al. 2008).

### WGS of the 1296 Zhejiang Mtb strains

Genomic DNA was sequenced using an Illumina HiSeq 2000 with an expected coverage of 100X. Paired-end reads were mapped to the reference genome, H37Rv (GenBank AL123456), using the Bowtie 2 software. The SAMtools (version 1.6)/BCFtools suite was used to call fixed SNPs (frequency ≥95%) (Li et al. 2009). We excluded all SNPs that were located in repetitive regions of the genome (e.g., PPE/PE/PGRS family genes, phage sequences, insertions and mobile genetic elements), as it is difficult to characterize such regions with short-read sequencing technologies (Yang et al. 2017). Small insertions or deletions, which were identified by VarScan (version 2.3.9) (Koboldt et al. 2012), were also excluded.

### Collection of the relevant WGS data

In order to construct phylogenetic trees including global strains and our samples, we curated a collection of MTBC representing geographic and genetic diversity. WGS data from global *Mycobacterium tuberculosis complex* (MTBC) lineage 2 and lineage 4 isolates was identified by searching PubMed for articles with WGS data. We downloaded the original sequencing reads from the European Nucleotide Archive (EMBL-EBI) and extracted the geographic origin and year of collection for each isolate from the relevant article. If the paper did not include this information, we sent an inquiry to the authors. Sequencing data were downloaded for 1154 MTBC isolates and geographic information was obtained for 1153 isolates (Supplementary_Table_S2).

### Phylogenetic analysis and pairwise determination of SNP distances

The fixed SNPs, excluding those in the proline-glutamic acid-proline-proline-glutamic acid sequence, the proline-glutamic acid-polymorphic GC-rich sequence and drug resistance-associated genes, were combined into a concatenated alignment. The best-scoring maximum likelihood phylogenetic tree was computed using RAxML v7.4.2 (Alexandros 2014) based on the concatenated alignment of 98,672 sites spanning the whole genome. Given the considerable size of the dataset (1296 Zhejiang strains + 161 of 1154 global strains from China + 21 reference strains (Zhang et al. 2013; Liu et al. 2018); 98,672 SNP sites), the rapid bootstrapping algorithm (*N*=100, *x*=12,345) and maximum likelihood search were used to construct the phylogenetic tree. The resulting tree was rooted on *M. canettii* (GenBank accession number: NC_019950.1). Lineage-defining nodes were based on 21 widely used isolates representing the six main phylogeographic lineages of MTBC. Bootstrap values were computed to assess the confidence of each clade, and to ensure that all lineage-defined nodes were highly supported (95-100%).

Filtered SNPs from isolates of lineages 2 and 4 were combined into a concatenated alignment as a fasta file. Pairwise SNP distances were calculated with the Bio:SeqIO package (Hackett et al. 2015). A pairwise SNP distance to all isolates of the same lineage was calculated for each isolate, and a distribution of the mean pairwise distance was plotted.

### Bayesian-based coalescent analysis

We randomly selected 197 Mtb strains from published studies (Zhang et al. 2013; Liu et al. 2018) to represent the national diversity (31 out of the 34 provincial regions of China) of Mtb sub-lineages in China and 48 Mtb strains from Zhejiang to represent the provincial diversity (collected from four regions [eastern/northern/western/southern Zhejiang] in 1998/2003/2008/2013, ignoring strains from middle Zhejiang to avoid confusion in constructing transmission routes) (Supplementary_Table_S3). The 197 and 48 strains were used for national and provincial phylogenetic reconstructions, respectively.

We applied Beast (Bayesian evolutionary analysis by sampling trees) (version 1.8.4) (Drummond et al. 2012), a genetic analysis software package based on the Monte Carlo Markov Chain algorithm (MCMC), to estimate the mutation rate, the divergence time of the Mtb strains and the times of the most recent common ancestors (tMRCAs) for lineages 2 and 4 and their sub-lineages. First, we imported the fasta file containing the genome sequencing information for the 197/48 strains into BEAUti software. To determine the Mtb genome substitution rate, we imposed a normal distribution for the substitution rate of Mtb with a mean of 4.6 × 10^−8^ substitutions per genome per site per year (95% highest posterior density [HPD] interval: 3.0 × 10^−8^ to 6.2 × 10^−8^), as described in a previous study (Bos et al. 2014). For the prior distribution of tMRCA, we imposed a normal distribution with a mean of 13,500 and a SE of 3000, as previously applied by Lin et al. (N 2014). We used an uncorrelated log-normal distribution for the substitution rate, an optimal evolution model of GTR+Γ4 (general time reversible + gamma-distributed rate variation with four rate categories), and the evolution model that was selected using Jmodeltest version 2.1.7.

To obtain reliable results, we ran a chain of 1 × 10^8^ generations, sampling every 10,000 generations to ensure independent convergence of the chain. Convergence was assessed using Tracer (version 1.7.0) (Liu et al. 2018), ensure that all relevant parameters reached an effective sample size of >200. The first 10% of the chain was discarded as burn-in, and we used the remaining chain to construct a Maximum Clade Credibility Tree (MCC tree) using Tree Annotator (version 1.8.4). Phylogenetic trees were visualized using FigTree (version 1.4.3). (Liu et al. 2018)

### Calculation of dN/dS ratios

To assess the antigenic diversity of human T cell epitopes among our Mtb samples, we chose a set of 491 epitopes corresponding to 130 non-overlapping regions in the antigen alignment (Comas et al. 2010). To assess how other regions of the genome are evolving, we also obtained alignments for essential and nonessential genes. Alignments of epitopes and non-epitope-containing regions for antigens, as well as essential and nonessential genes, were used to calculate pairwise dN/dS ratios for lineages 2 and 4. Pairwise dN and dS values within each lineage were calculated using the R package tool, seqinr, with the ka/ks function (Comas et al. 2010). To avoid having undetermined pairwise dN/dS values due to dN or dS being zero, we calculated a mean dN/dS value for each sequenced isolate by dividing its mean pairwise dN by its mean pairwise dS with respect to all other sequenced isolates within each lineage.

## Data Availability Statement

The data underlying this article will be shared on reasonable request to the corresponding author.

## Acknowledgements

We gratefully acknowledge our funders.

## Author Contributions

BW, YW, QW, XW and WW designed the study. BW, LZ, ZL, SC and XW collected and contributed the MTBC isolates analysed in this study. YW, QW, MB and WW analysed the sequencing reads and performed the genetic analysis. YW, LC and LB participated in the analysis of integrating tuberculosis history with Chinese human population history. QW and WZ performed the statistical analysis. BW, YW, QW, XW and WW drafted the manuscript. MB, BK revised the structure of this paper and polished the language. All authors critically reviewed and approved the final version of the manuscript.

## References

Alexandros S. 2014. RAxML version 8: a tool for phylogenetic analysis and post-analysis of large phylogenies. Bioinformatics 30: 1312–1313

Aziz, M.A, Wright, Muynck D, Laszlo. 2008. Anti-tuberculosis drug resistance in the world - third global report. Geneva, Switzerland, World Health Organization [WHO], 2008 12: 257–261

Barton L, Newsome SD, Chen FH, Wang H, Guilderson TP, Bettinger RL. 2009. Agricultural origins and the isotopic identity of domestication in northern China. Proc Natl Acad Sci USA 106: 5523–5528.doi:10.1073/pnas.0809960106

Bos KI, Harkins KM, Herbig A, Coscolla M, Weber N, Comas I, Forrest SA, Bryant JM, Harris SR, Schuenemann VJ, Campbell TJ, Majander K, Wilbur AK, Guichon RA, Wolfe Steadman DL, Cook DC, Niemann S, Behr MA, Zumarraga M, Bastida R, Huson D, Nieselt K, Young D, Parkhill J, Buikstra JE, Gagneux S, Stone AC, Krause J. 2014. Pre-Columbian mycobacterial genomes reveal seals as a source of New World human tuberculosis. Nature 514: 494–497.doi:10.1038/nature13591

CCTV. 2007. What is the “Silk Road” and “Maritime Silk Road”?,

China NHaFPCotPsRo. 2017. Diagnosis for pulmonary tuberculosis. Health Industry Standards of the People’s Republic of China

Coll F, McNerney R, Guerra-Assuncao JA, Glynn JR, Perdigao J, Viveiros M, Portugal I, Pain A, Martin N, Clark TG. 2014. A robust SNP barcode for typing Mycobacterium tuberculosis complex strains. Nat Commun 5: 4812.doi:10.1038/ncomms5812

Comas I, Chakravartti J, Small PM, Galagan J, Niemann S, Kremer K, Ernst JD, Gagneux S. 2010. Human T cell epitopes of Mycobacterium tuberculosis are evolutionarily hyperconserved. Nat Genet 42: 498–503.doi:10.1038/ng.590

Comas I, Coscolla M, Luo T, Borrell S, Holt KE, Kato-Maeda M, Parkhill J, Malla B, Berg S, Thwaites G, Yeboah-Manu D, Bothamley G, Mei J, Wei L, Bentley S, Harris SR, Niemann S, Diel R, Aseffa A, Gao Q, Young D, Gagneux S. 2013. Out-of-Africa migration and Neolithic coexpansion of Mycobacterium tuberculosis with modern humans. Nat Genet 45: 1176–1182.doi:10.1038/ng.2744

Coscolla M, Copin R, Sutherland J, Gehre F, de Jong B, Owolabi O, Mbayo G, Giardina F, Ernst JD, Gagneux S. 2015. M. tuberculosis T Cell Epitope Analysis Reveals Paucity of Antigenic Variation and Identifies Rare Variable TB Antigens. Cell Host Microbe 18: 538–548.doi:10.1016/j.chom.2015.10.008

Coscolla M, Gagneux S. 2010. Does M. tuberculosis genomic diversity explain disease diversity? Drug Discov Today Dis Mech 7: e43–e59.doi:10.1016/j.ddmec.2010.09.004

Coscolla M, Gagneux S. 2014. Consequences of genomic diversity in Mycobacterium tuberculosis. Semin Immunol 26: 431–444.doi:10.1016/j.smim.2014.09.012

Drummond AJ, Suchard MA, Xie D, Rambaut A. 2012. Bayesian phylogenetics with BEAUti and the BEAST 1.7. Mol Biol Evol 29: 1969–1973.doi:10.1093/molbev/mss075

Ernst JD. 2012. The immunological life cycle of tuberculosis. Nat Rev Immunol 12: 581–591.doi:10.1038/nri3259

Fan JJ. 2017. The pictures of the Battle of Mingtiao. China surveying and mapping: 61–63

Filliol I, Motiwala AS, Cavatore M, Qi W, Hazbon MH, Bobadilla del Valle M, Fyfe J, Garcia-Garcia L, Rastogi N, Sola C, Zozio T, Guerrero MI, Leon CI, Crabtree J, Angiuoli S, Eisenach KD, Durmaz R, Joloba ML, Rendon A, Sifuentes-Osornio J, Ponce de Leon A, Cave MD, Fleischmann R, Whittam TS, Alland D. 2006. Global phylogeny of Mycobacterium tuberculosis based on single nucleotide polymorphism (SNP) analysis: insights into tuberculosis evolution, phylogenetic accuracy of other DNA fingerprinting systems, and recommendations for a minimal standard SNP set. J Bacteriol 188: 759–772.doi:10.1128/JB.188.2.759-772.2006

Ford CB, Lin PL, Chase MR, Shah RR, Iartchouk O, Galagan J, Mohaideen N, Ioerger TR, Sacchettini JC, Lipsitch M, Flynn JL, Fortune SM. 2011. Use of whole genome sequencing to estimate the mutation rate of Mycobacterium tuberculosis during latent infection. Nat Genet 43: 482–486.doi:10.1038/ng.811

Gagneux S. 2012. Host-pathogen coevolution in human tuberculosis. Philos Trans R Soc Lond B Biol Sci 367: 850–859.doi:10.1098/rstb.2011.0316

Glaziou P, Falzon D, Floyd K, Raviglione M. 2013. Global epidemiology of tuberculosis. Semin Respir Crit Care Med 34: 3–16.doi:10.1055/s-0032-1333467

Gray RD, Jordan FM. 2000. Language trees support the express-train sequence of Austronesian expansion. Nature 405: 1052–1055.doi:10.1038/35016575

Hackett R, Moulton OC, Raff JW. 2015. Biology Open: evaluating impact. Biology Open 4: 1609

Hershberg R, Lipatov M, Small PM, Sheffer H, Niemann S, Homolka S, Roach JC, Kremer K, Petrov DA, Feldman MW, Gagneux S. 2008. High functional diversity in Mycobacterium tuberculosis driven by genetic drift and human demography. PLoS Biol 6: e311.doi:10.1371/journal.pbio.0060311

Koboldt DC, Zhang Q, Larson DE, Shen D, McLellan MD, Lin L, Miller CA, Mardis ER, Ding L, Wilson RK. 2012. VarScan 2: somatic mutation and copy number alteration discovery in cancer by exome sequencing. Genome Res 22: 568–576.doi:10.1101/gr.129684.111

Leung CC, Chee CBE, Zhang Y. 2018. Tuberculosis updates 2018: Innovations and developments to end TB. Respirology 23: 356–358.doi:10.1111/resp.13244

Li H, Handsaker B, Wysoker A, Fennell T, Ruan J, Homer N, Marth G, Abecasis G, Durbin R, Genome Project Data Processing S. 2009. The Sequence Alignment/Map format and SAMtools. Bioinformatics 25: 2078–2079.doi:10.1093/bioinformatics/btp352

Li QHW, W. Z. 1981. The picture of Qin conquering the other six countries. History Teaching: 65

Liu Q, Ma A, Wei L, Pang Y, Wu B, Luo T, Zhou Y, Zheng HX, Jiang Q, Gan M, Zuo T, Liu M, Yang C, Jin L, Comas I, Gagneux S, Zhao Y, Pepperell CS, Gao Q. 2018. China’s tuberculosis epidemic stems from historical expansion of four strains of Mycobacterium tuberculosis. Nat Ecol Evol 2: 1982–1992.doi:10.1038/s41559-018-0680-6

Lonnroth K, Jaramillo E, Williams BG, Dye C, Raviglione M. 2009. Drivers of tuberculosis epidemics: the role of risk factors and social determinants. Soc Sci Med 68: 2240–2246.doi:10.1016/j.socscimed.2009.03.041

Mokrousov I. 2013. Insights into the origin, emergence, and current spread of a successful Russian clone of Mycobacterium tuberculosis. Clin Microbiol Rev 26: 342–360.doi:10.1128/CMR.00087-12

N L. 2014. Genome-wide analysis of the populations of Mycobacterium tuberculosis from China.

Nathanson E, Nunn P, Uplekar M, Floyd K, Jaramillo E, Lonnroth K, Weil D, Raviglione M. 2010. MDR tuberculosis--critical steps for prevention and control. N Engl J Med 363: 1050–1058.doi:10.1056/NEJMra0908076

Oleksyk TK, Smith MW, O’Brien SJ. 2010. Genome-wide scans for footprints of natural selection. Philos Trans R Soc Lond B Biol Sci 365: 185–205.doi:10.1098/rstb.2009.0219

Parwati I, van Crevel R, van Soolingen D. 2010. Possible underlying mechanisms for successful emergence of the Mycobacterium tuberculosis Beijing genotype strains. Lancet Infect Dis 10: 103–111.doi:10.1016/S1473-3099(09)70330-5

Reed MB, Pichler VK, McIntosh F, Mattia A, Fallow A, Masala S, Domenech P, Zwerling A, Thibert L, Menzies D, Schwartzman K, Behr MA. 2009. Major Mycobacterium tuberculosis lineages associate with patient country of origin. J Clin Microbiol 47: 1119–1128.doi:10.1128/JCM.02142-08

Rodrigo T, Cayla JA, Garcia de Olalla P, Galdos-Tanguis H, Jansa JM, Miranda P, Brugal T. 1997. Characteristics of tuberculosis patients who generate secondary cases. Int J Tuberc Lung Dis 1: 352–357

Shih CH, Chang CM, Lin YS, Lo WC, Hwang JK. 2012. Evolutionary information hidden in a single protein structure. Proteins 80: 1647–1657.doi:10.1002/prot.24058

Stucki D, Brites D, Jeljeli L, Coscolla M, Liu Q, Trauner A, Fenner L, Rutaihwa L, Borrell S, Luo T, Gao Q, Kato-Maeda M, Ballif M, Egger M, Macedo R, Mardassi H, Moreno M, Tudo Vilanova G, Fyfe J, Globan M, Thomas J, Jamieson F, Guthrie JL, Asante-Poku A, Yeboah-Manu D, Wampande E, Ssengooba W, Joloba M, Henry Boom W, Basu I, Bower J, Saraiva M, Vaconcellos SEG, Suffys P, Koch A, Wilkinson R, Gail-Bekker L, Malla B, Ley SD, Beck HP, de Jong BC, Toit K, Sanchez-Padilla E, Bonnet M, Gil-Brusola A, Frank M, Penlap Beng VN, Eisenach K, Alani I, Wangui Ndung’u P, Revathi G, Gehre F, Akter S, Ntoumi F, Stewart-Isherwood L, Ntinginya NE, Rachow A, Hoelscher M, Cirillo DM, Skenders G, Hoffner S, Bakonyte D, Stakenas P, Diel R, Crudu V, Moldovan O, Al-Hajoj S, Otero L, Barletta F, Jane Carter E, Diero L, Supply P, Comas I, Niemann S, Gagneux S. 2016. Mycobacterium tuberculosis lineage 4 comprises globally distributed and geographically restricted sublineages. Nat Genet 48: 1535–1543.doi:10.1038/ng.3704

Subbian S, Bandyopadhyay N, Tsenova L, O’Brien P, Khetani V, Kushner NL, Peixoto B, Soteropoulos P, Bader JS, Karakousis PC, Fallows D, Kaplan G. 2013. Early innate immunity determines outcome of Mycobacterium tuberculosis pulmonary infection in rabbits. Cell Commun Signal 11: 60.doi:10.1186/1478-811X-11-60

Tsenova L, Ellison E, Harbacheuski R, Moreira AL, Kurepina N, Reed MB, Mathema B, Barry CE, 3rd, Kaplan G. 2005. Virulence of selected Mycobacterium tuberculosis clinical isolates in the rabbit model of meningitis is dependent on phenolic glycolipid produced by the bacilli. J Infect Dis 192: 98–106.doi:10.1086/430614

Tuite AR, Guthrie JL, Alexander DC, Whelan MS, Lee B, Lam K, Ma J, Fisman DN, Jamieson FB. 2013. Epidemiological evaluation of spatiotemporal and genotypic clustering of Mycobacterium tuberculosis in Ontario, Canada. Int J Tuberc Lung Dis 17: 1322–1327.doi:10.5588/ijtld.13.0145

Walker TM, Ip CL, Harrell RH, Evans JT, Kapatai G, Dedicoat MJ, Eyre DW, Wilson DJ, Hawkey PM, Crook DW, Parkhill J, Harris D, Walker AS, Bowden R, Monk P, Smith EG, Peto TE. 2013. Whole-genome sequencing to delineate Mycobacterium tuberculosis outbreaks: a retrospective observational study. Lancet Infect Dis 13: 137–146.doi:10.1016/S1473-3099(12)70277-3

WHO. 2018. WHO global tuberculosis control report 2018..

Wirth T, Hildebrand F, Allix-Beguec C, Wolbeling F, Kubica T, Kremer K, van Soolingen D, Rusch-Gerdes S, Locht C, Brisse S, Meyer A, Supply P, Niemann S. 2008. Origin, spread and demography of the Mycobacterium tuberculosis complex. PLoS Pathog 4: e1000160.doi:10.1371/journal.ppat.1000160

Yang C, Luo T, Shen X, Wu J, Gan M, Xu P, Wu Z, Lin S, Tian J, Liu Q. 2017. Transmission of multidrug-resistant Mycobacterium tuberculosis in Shanghai, China: a retrospective observational study using whole-genome sequencing and epidemiological investigation. Lancet Infectious Diseases 17: 275–284

Yi H. 2019. Liangzhu Culture and Huaxia Civilization. The Central Plains Culture Research 7: 5–13.doi:10.16600/j.cnki.41-1426/c.2019.05.001

Yruela I, Contreras-Moreira B, Magalhaes C, Osorio NS, Gonzalo-Asensio J. 2016. Mycobacterium tuberculosis Complex Exhibits Lineage-Specific Variations Affecting Protein Ductility and Epitope Recognition. Genome Biol Evol 8: 3751–3764.doi:10.1093/gbe/evw279

Zhang H, Li D, Zhao L, Fleming J, Lin N, Wang T, Liu Z, Li C, Galwey N, Deng J, Zhou Y, Zhu Y, Gao Y, Wang T, Wang S, Huang Y, Wang M, Zhong Q, Zhou L, Chen T, Zhou J, Yang R, Zhu G, Hang H, Zhang J, Li F, Wan K, Wang J, Zhang XE, Bi L. 2013. Genome sequencing of 161 Mycobacterium tuberculosis isolates from China identifies genes and intergenic regions associated with drug resistance. Nat Genet 45: 1255–1260.doi:10.1038/ng.2735

Zhong H, Shi H, Qi XB, Duan ZY, Tan PP, Jin L, Su B, Ma RZ. 2011. Extended Y chromosome investigation suggests postglacial migrations of modern humans into East Asia via the northern route. Mol Biol Evol 28: 717–727.doi:10.1093/molbev/msq247

