## Supplemental Materials for "Genetic composition and evolution of the prevalent *Mycobacterium* tuberculosis lineages 2 and 4 in the Chinese and Zhejiang Province populations"

**This PDF file includes:**

Figure S1 to Figure S5

**Other supplementary materials for this manuscript include the following:**

Table S1 to Table S3 (Excel format files)

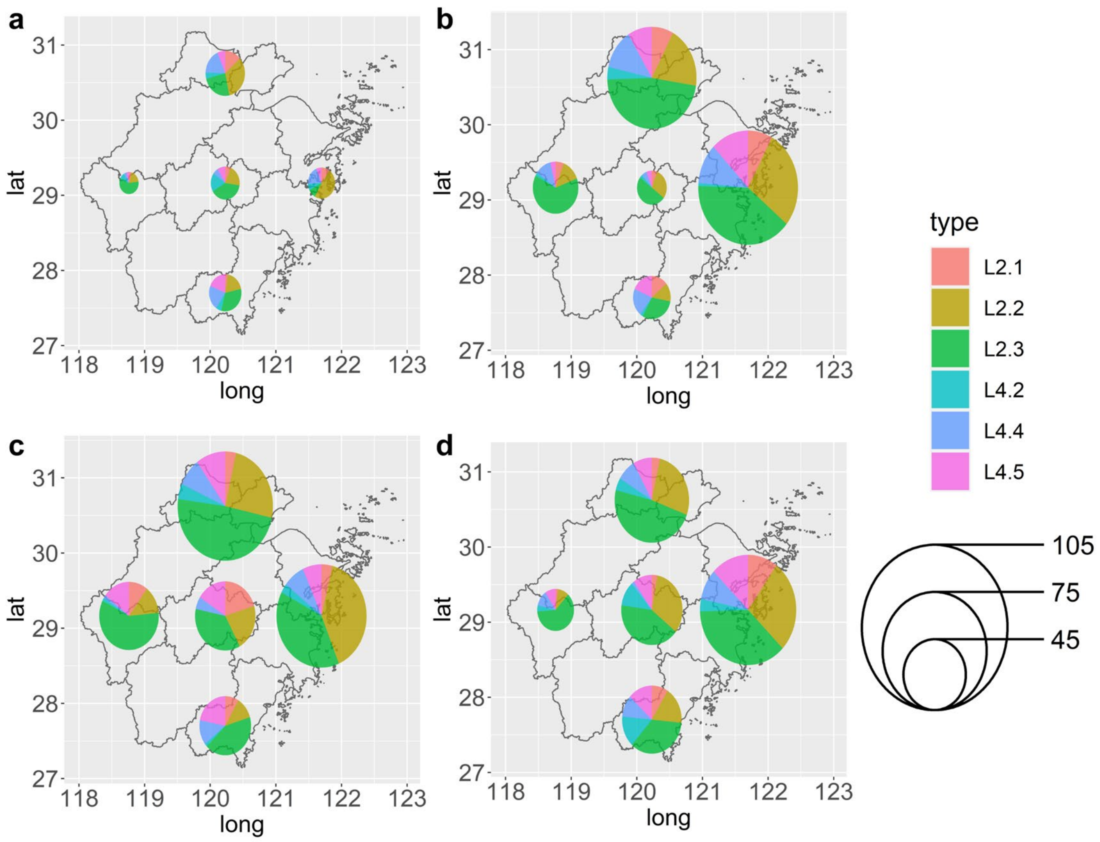


**Figure S1.** Changes of the distribution of Mycobacterium tuberculosis sub-lineages in Zhejiang Province from 1998 to 2013. a) 1998, b) 2003, c) 2008, d) 2013. The size of the circle represents the sample size of the strains.

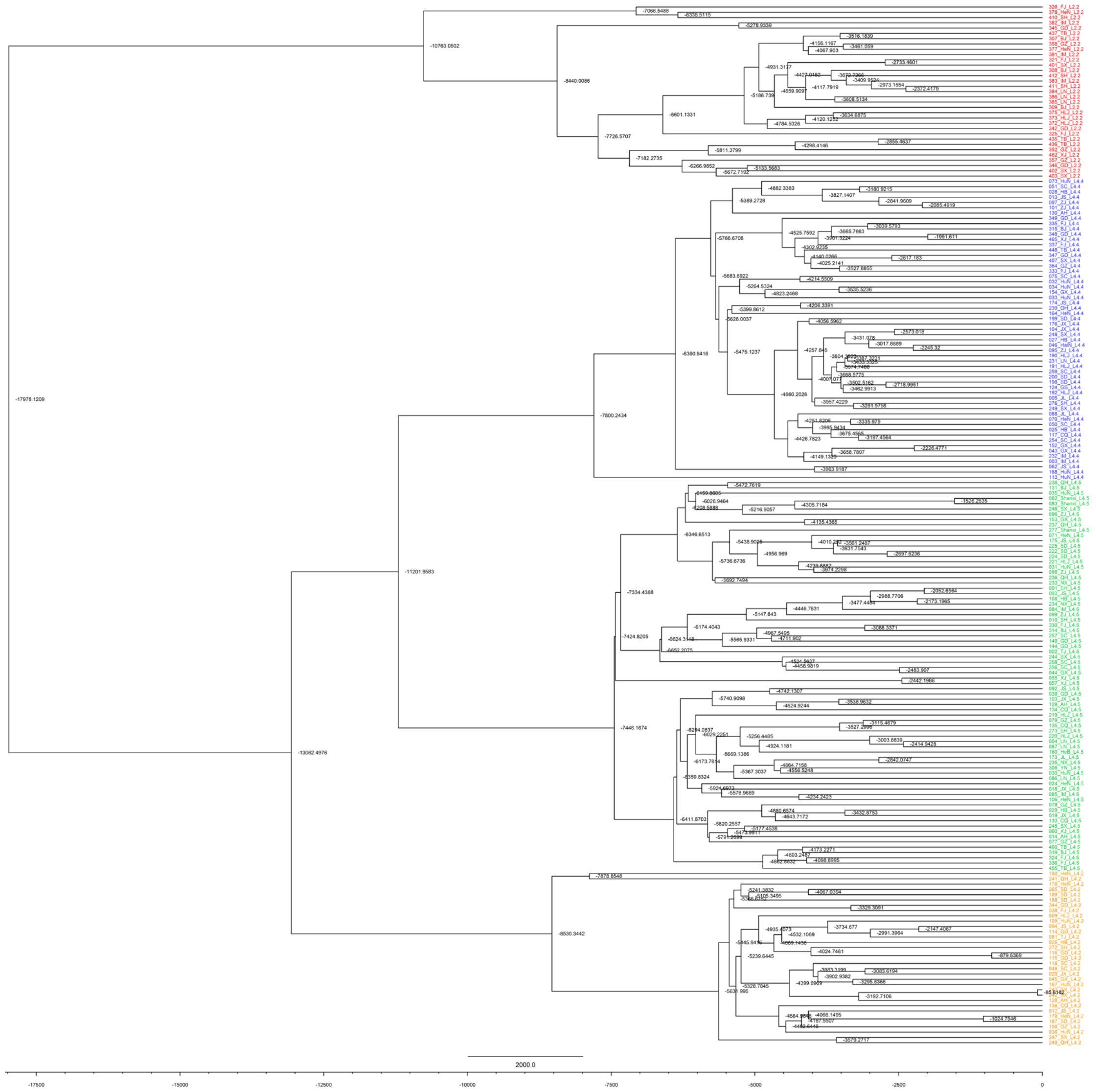


**Figure S2.** Phylogenetic tree of Mycobacterium tuberculosis in China.

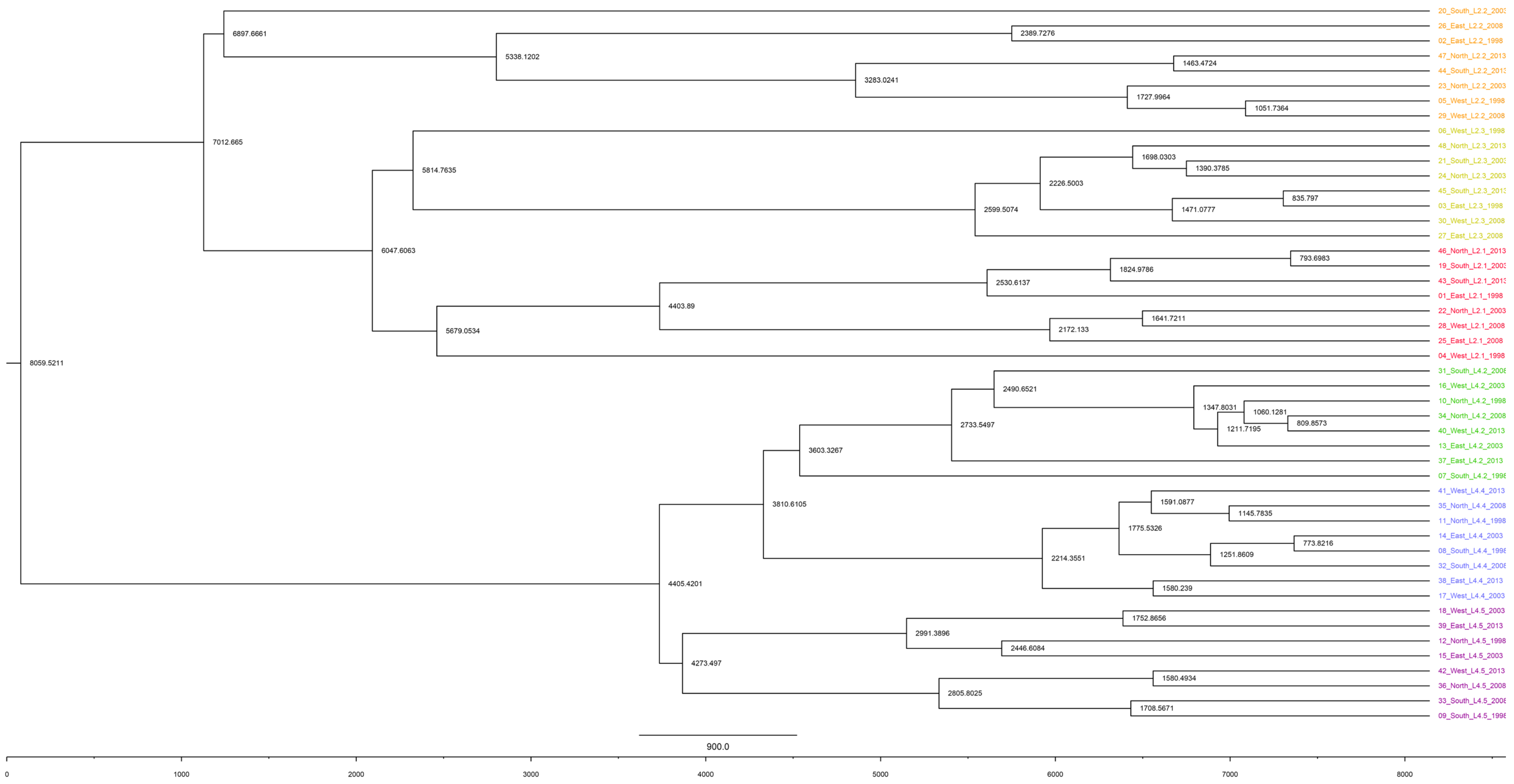

**Figure S3.** Phylogenetic tree of Mycobacterium tuberculosis in Zhejiang Province.

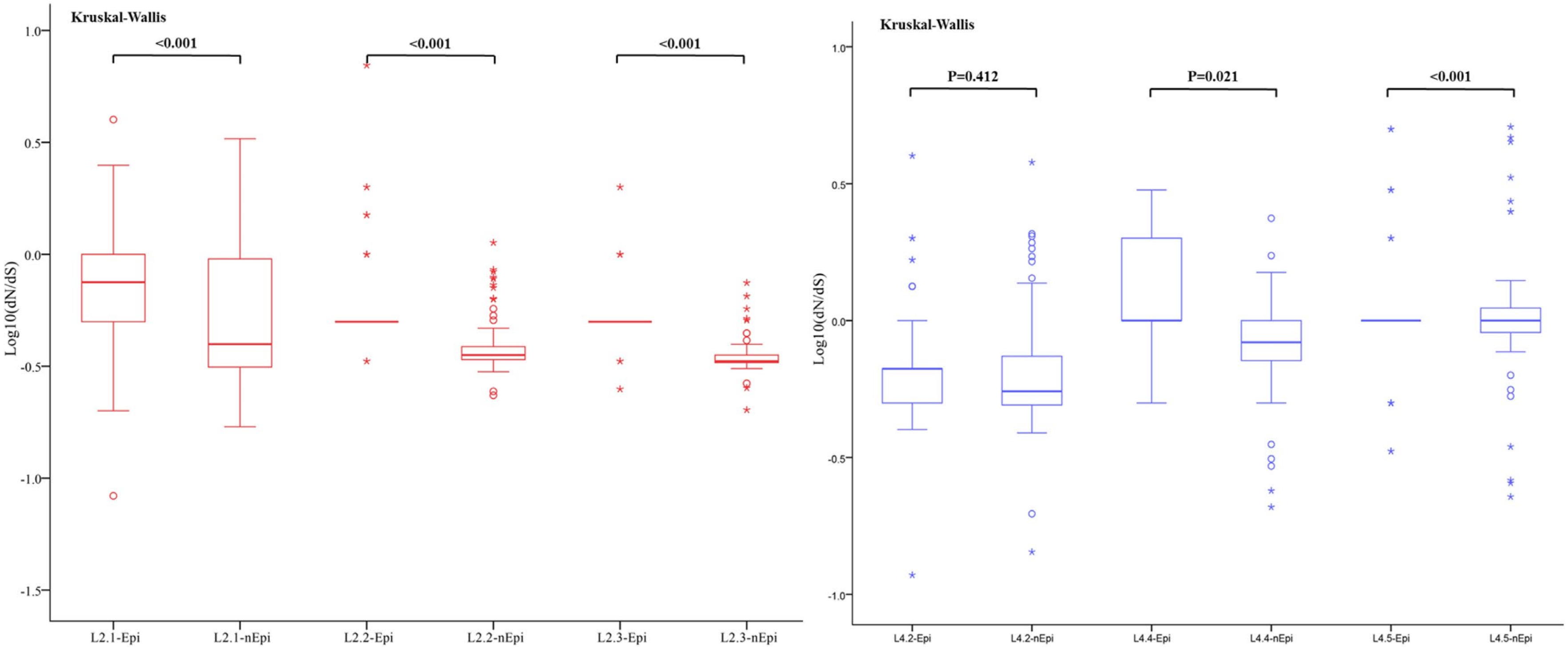


**Figure S4.** Pairwise ratios of rates of nonsynonymous to synonymous substitutions (dN/dS) in sublineages in lineage 2 and lineage 4 for epitopes and non-epitope regions of T cell antigens.

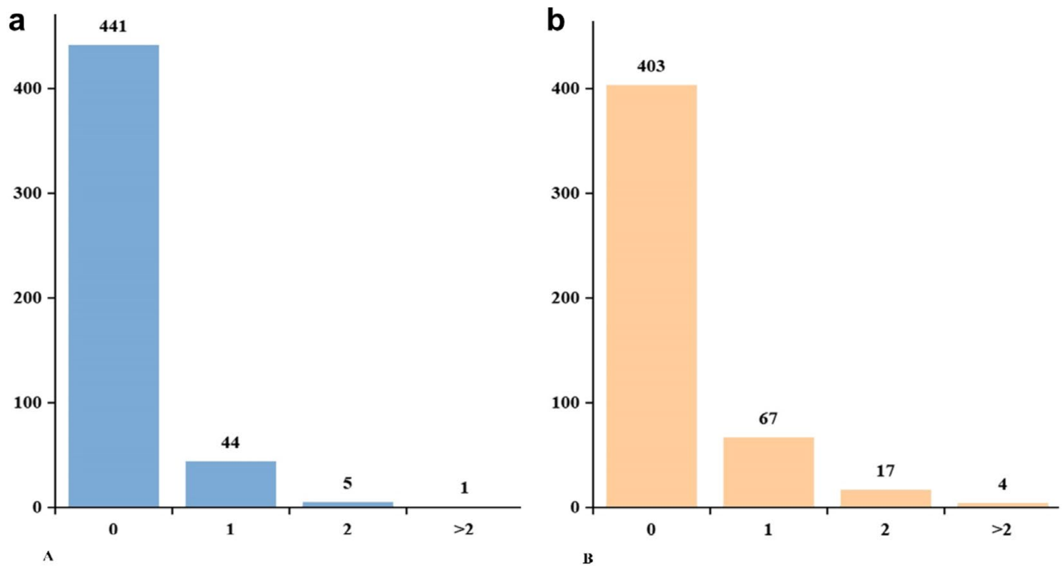


**Figure S5.** Frequency distribution of the number of epitopes with nonsynonymous variants. A total of 491 T cell epitopes were included in the analysis. The number above each bar corresponds to the epitope count. a) lineage 4, b) lineage 2.
